# Novel GC–MS/MS Strategy for Fructose Quantification and Stable Isotope Tracing: Development, Validation, and SIM vs MRM Comparison

**DOI:** 10.64898/2026.08.24.746767

**Authors:** Melany Rios-Morales, Florine H.M. Westerbeke, Max Nieuwdorp, Frédéric M. Vaz, Dewi van Harskamp

**Affiliations:** Amsterdam UMC, Department of Experimental Vascular Medicine, Amsterdam, the Netherlands; Amsterdam UMC, Department of Internal and Vascular Medicine, Amsterdam, the Netherlands; Amsterdam UMC, Department of Laboratory Medicine, Laboratory Genetic Metabolic Diseases, Emma Children’s Hospital, Amsterdam, The Netherlands; Amsterdam UMC, Amsterdam Gastroenterology Endocrinology Metabolism, Inborn errors of metabolism, Amsterdam, The Netherlands; Amsterdam UMC, Core Facility Metabolomics, Amsterdam, The Netherlands; Amsterdam UMC, Emma Center for Personalized Medicine, Amsterdam, Netherlands

**Keywords:** Fructose, gas chromatography, mass spectrometry, stable isotope

## Abstract

High dietary fructose consumption is a major contributor to the development of obesity and related cardiometabolic diseases, highlighting the need for accurate assessment of fructose metabolism in humans. Stable isotope tracer approaches, such as ^13^C_6_-fructose, require highly sensitive and specific analytical methods to quantify both concentrations and isotopic enrichments.

In this study, we developed and validated a robust gas chromatography–triple quadrupole mass spectrometry (GC– QQQ)-based method for the simultaneous measurement of unlabeled and ^13^C_6_-fructose in human plasma. The method employs oximation and per-acetate derivatization, and demonstrates high specificity and accuracy. Intra- and inter-assay precision were below 10%, with no detectable carry-over, and a lower limit of quantification (LLOQ) of 0.1 nmol/mL for concentration and 0.02 molar percent excess (MPE%) for enrichment and no interference from glucose.

We further compared data acquisition using multiple reaction monitoring (MRM) and selected ion monitoring (SIM). MRM showed superior performance at the low concentrations and enrichment levels characteristic of clinical plasma samples, resulting in improved sensitivity and lower LLOQs compared to SIM.

Overall, this validated method provides a sensitive and reliable approach for fructose tracer studies in humans. Its application will facilitate robust investigations into fructose metabolism, and its role in metabolic dysregulation and obesity-related disease.

## Introduction

Obesity has emerged as one of the most significant global public health challenges, affecting both children and adults. In children aged 5–14 years, global obesity prevalence rose from 2.0% in 1990 to 6.9% in 2021, and is projected to reach 15.6% by 2050 Kerr, Patton (1). Similarly, the number of adults living with overweight and obesity exceeded 2 billion individuals worldwide in 2021 and is expected to approach 3.8 billion by 2050, likely representing more than half of the global adult population (2). This growing epidemic places an enormous burden on healthcare systems through increased rates of related cardiometabolic diseases including type 2 diabetes (T2D) and metabolic dysfunction-associated steatotic liver disease (MASLD). The economic burden of obesity is reflected in increasing healthcare costs and reduced productivity caused by illness and disability. Analyses suggest that if current trends persist, the global cost of overweight and obesity could reach about $4.32 trillion per year by 2035 (3).

The global rise in obesity is widely attributed to a chronic imbalance between energy intake and energy expenditure, though alternative theories suggest the process is more complex (4-6). One important example is the carbohydrate–insulin model, which proposes that diet composition plays a central role in driving fat storage and appetite regulation, contributing to dysregulated energy balance and increased hunger (5-7). Among the dietary carbohydrates implicated in this dysregulation, fructose has received particular attention due to its increased consumption in modern diets and its unique catabolic pathway (8-15).

Fructose is classically described as being metabolized in the liver into fatty acids through an unregulated pathway, leading to high circulating lipids levels and promoting hepatic steatosis (16-18). Furthermore, the rapid hepatic phosphorylation of fructose to fructose-1-phosphate can deplete hepatic ATP, triggering uric acid production and oxidative stress, which further exacerbate insulin resistance and de novo lipogenesis (DNL) (19-26). However, emerging pre-clinical evidence challenges the view of the liver as the exclusive site of fructose catabolism (27), positioning the small intestine as a key catabolic organ. At physiological doses, the intestine could convert fructose into glucose and organic acids to protect the liver from direct exposure; however, this protective capacity saturates at higher fructose loads, allowing excess fructose to reach both the liver and the distal intestine. In the latter, gut microbiota can ferment fructose into lipogenic substrates such as acetate and ethanol (28-31). This dose-dependent shift in fructose handling underscores the importance of accurately quantifying fructose to understand its role in metabolic disease.

Many of the proposed mechanisms underlying fructose-induced metabolic dysfunction have been established primarily in *in vitro* and animal models, thus, human studies are needed to fully understand fructose catabolism and its metabolic effects. Stable isotope tracer studies provide a useful approach to investigate metabolic pathways and substrate fate *in vivo* (32). However, accurate quantification of fructose kinetics is analytically challenging due to the relatively low concentrations of fructose in plasma compared with glucose (33, 34). To saturate all proposed processes and reach peripheral circulation, a high fructose load is needed. Due to the high price of isotopically labeled tracers, the tracer is only supplied as a small percentage of the total fructose load, resulting in low tracer enrichments, further increasing the need for a highly sensitive analytical method. We have previously shown that gas chromatography–triple quadrupole mass spectrometry (GC-QQQ) operated in multiple reaction monitoring (MRM) mode enables highly sensitive and precise measurement of isotopic enrichments (35). The aim of this study was to develop and validate an improved method for the analysis of fructose enrichments, ultimately facilitating robust human studies that will further elucidate the role of fructose catabolism in metabolic dysregulation and obesity-related disease. Specifically, we aimed to investigate whether oximation and per-acetate derivatization, commonly used for glucose analysis, is also suitable for fructose measurement using GC-QQQ, and whether the improved performance of MRM compared with selected ion monitoring (SIM) justifies the use of a more expensive analytical setup. Here, we present a comprehensive analytical validation of fructose measurement using this approach.

## Experimental

### Reagents

Methanol, hexane, dichloromethane, pyridine and ethyl acetate were purchased from BioSolve (the Netherlands). Hydroxylamine HCl was obtained from Sigma (Merck, Germany), and acetic anhydride from Alfa Aesar (Thermo, USA). To prevent spoiling of the acetic anhydride, the bottle was flushed with nitrogen after each use.

### Standard preparation

Stock solutions of 10 mM fructose (Sigma, Merck, Germany), 10 mM D_2_-fructose (Campro, the Netherlands) and 10 mM ^13^C_6_-fructose (Cambridge Isotope Laboratories, USA) were prepared in 20% methanol in water. Aliquots were stored at -20°C, and a calibration curve was prepared fresh for every analysis. The calibration curve ranged from 0 to 1000 nmol/ml for concentration, and from 0 to 1.5 molar percent excess (MPE%) for enrichment. The calibration curves for ^13^C_6_-fructose enrichment and fructose concentration (using D_2_-fructose as internal standard (IS)) were established within a single standard line, yielding linear results; however, to avoid artificial correlation, the concentration and enrichment values were deliberately arranged in a non-sequential and mixed manner.

### Control samples

Control samples were prepared from pooled plasma (purchased from Sanquin, the Netherlands) spiked at different levels with fructose and ^13^C_6_-fructose. Three pools were used: a blank pool (unspiked, but with endogenous fructose present), low, spiked with approximately 100 nmol/ml unlabeled fructose and 0.5 MPE% ^13^C_6_-fructose, spiked with approximately 600 nmol/ml unlabeled fructose and 1.5 MPE% ^13^C_6_-fructose. The control samples were stored at -20°C and thawed before use.

### Clinical samples

Human plasma samples for fructose measurement were obtained from the Fecal microbiota transplantation for improved Fructose catabolism and Insulin sensiTivity In patients with metabolic syndrome (FFIT) trial (Dutch Clinical Trial Register, registration ID: NL-OMON24239, registered on 18-02-2022). Briefly, after an overnight fast subjects performed a fructose challenge test (FCT) where they ingested 1 g/kg bodyweight of unlabeled fructose dissolved in water followed by a capsule containing 120 mg of ^13^C_6_-fructose as a tracer. Blood samples were collected at time = 0, 30, 60, 90, 120, 150, 180, 240, and 360 min to characterize fructose appearance in peripheral circulation. In addition, at each time point, plasma collected in heparin and EDTA tubes was compared to evaluate the impact of anticoagulant type on fructose measurement, providing guidance for future clinical studies.

### Preparation

In an Eppendorf tube, 100 µl sample, 10 µl of 1 mM D_2_-fructose and 900 µl methanol were added. The samples were incubated on ice for 10 minutes, before centrifugation for 5 minutes at 2500 rpm at 4°C. All supernatant was transferred (∼1ml) into a clean glass tube. Then, 1 ml of hexane was added, vortexed for 1 minute at 1500 rpm and the samples were centrifuged for 3 minutes at 2500 rpm, to wash away the lipids in the sample. The bottom polar layer was transferred into a clean 1.5 ml GC-vial. The extracts were dried under a gentle stream of nitrogen at 50°C. To remove trace amounts of water, 200 µl of dichloromethane was added to the sample, vortexed and dried under nitrogen at 50°C.

### Derivatization

To each sample, 100 µl of hydroxylamine solution was added (20 mg/ml hydroxylamine in pyridine, prepared fresh each analysis). The sample was then incubated for 30 minutes at 90°C, allowed to cool down to room temperature and dried under nitrogen at 50°C. Then, 200 µl acetic anhydride was added to the vial. This was incubated again for 30 minutes at 90°C. After cooling to room temperature, 500 µl of doubly distilled water and 500 µl of dichloromethane was added to the vials, vortexed for 1 minute, and centrifuged for 5 minutes at 2000 rpm. Six hundred µl of the bottom layer was transferred to a new vial, and dried under nitrogen at 50°C. The sample was then dissolved in 50 µl of ethyl acetate and transferred to an insert.

### Instrument

A 7890A GC, equipped with a 7693 auto sampler coupled to a 7000 Triple Quadrupole Mass Spectrometer (Agilent Technologies, USA), was used to perform all analyses. The GC was equipped with a Dean’s switch in order to reduce contamination of the source by sending the effluent prior to and after elution of peaks of interest to the waste. Samples were introduced through a temperature-programmed multimode inlet. The GC was fitted with a VF5ms column (30 m, 0.25 mm, 0.25 μm). Through the Dean’s switch, the flow was either directed to the mass spectrometer (uncoated fused silica capillary, 1.8 m, 0.180 mm) or to waste (uncoated fused silica capillary, 1.3 m, 0.180 mm). Helium 5.0 was used as carrier gas at a flow rate of 1.2 ml/min. One microliter of sample was injected per run, and each sample was analyzed in duplicate. The initial oven temperature was 55 °C for 1 min. The temperature was then increased by 30 °C/min to 160 °C and held constant for 0.5 min. It was then increased by 6 °C/min to 225 °C and held constant for 7 min, followed by an increase in temperature of 30°C/min to 300°C, and held constant for 10 min. The run time was 35.33 min. The system was operated in either the MRM mode or SIM mode, while recording the effluent from 20 to 24 min. The precursor ions were selected at a narrower resolution (target peak width of 0.45, in order to prevent a contribution from the neighboring mass to charge ratio (m/z) value to the m/z value of interest), and the product ions were selected with normal resolution (0.7). For SIM analysis, the resolution was set to the narrower resolution (0.45). All MRM transitions were recorded with a gain factor of 25, and SIM transitions with a gain factor of 10. The gain factors differ due to the upper limit of the linear detection range. SIM yields a higher signal since, unlike other modes, it omits fragmentation as part of its distinct measurement principle. The transitions were recorded with a dwell time of 150 ms. Collision Energy in MRM mode was 3 eV. Further MRM and SIM settings are shown in Table 1. The ion source installed was an Electron Ionization Extractor Ion Source, operated at 230 °C and 70 eV. The collision gas was nitrogen 6.0 at a flow rate of 1.5 ml/min, and the quench gas was helium at a flow rate of 2.25 ml/min. The transfer line temperature was kept at 280 °C, the temperatures of quadrupole 1 and 2 were kept at 150 °C. The acquisition software for operating the GC-QQQ was MassHunter Workstation GC/MS Data Acquisition Version 10.2 Service Release 3 Build 10.2.530.3 (Agilent Technologies, USA).

**Table 1.** Mass spectrometry conditions, Multiple reaction monitoring (MRM) and Single Ion Monitoring (SIM) settings for the fructose derivative (MW, molecular weight; m/z, mass to charge ratio)

| Mode | Isotope | MW derivative | Precursor ion ($m/z$) | Product ion ($m/z$) |
| --- | --- | --- | --- | --- |
| MRM | M | 387 | 345.1 | 183 |
|  | M+2 | 389 | 347.1 | 185 |
|  | M+6 | 393 | 351.1 | 189 |
| SIM | M | 387 | 345.1 | n.a. |
|  | M+2 | 389 | 347.1 | n.a. |
|  | M+6 | 393 | 351.1 | n.a. |

### Data analysis

Integration of the peak area was performed using MassHunter Workstation Quantitative Analysis for QQQ software Version 12.1 Build 12.1.938.3 (Agilent Technologies, USA) and subsequently converted into %MPE and concentrations (nmol/ml) by applying corrections based on the calibration curves. The slope and intercept of both the concentration and the enrichment curve were determined by linear regression analysis. Further calculations were performed using Excel 2016 (Microsoft Corporation, USA). Carryover was assessed using the statistical module specifically designed for carryover on the EP Evaluator software (Data Innovations LLC, Colchester, USA, Build 12.3.0.2).

### Method validation

#### Accuracy

To assess the accuracy of the method, the control samples were measured repeatedly. As pool plasma contains endogenous fructose, the unspiked (blank) pool was analyzed as well, and the increase in concentration and enrichment in the spiked pools were compared to the theoretical concentrations and enrichments. An accuracy of 100% ± 20% was considered acceptable.

#### Precision

Intra-assay and inter-assay precision for both enrichment and concentration were evaluated using the control pools. Intra-assay precision was determined from ten independently prepared replicates analyzed within a single day, while inter-assay precision was assessed from two aliquots per control pool, prepared on each of five distinct days. Each replicate was analyzed in duplicate. Coefficients of variation (CV%) were calculated from the resulting data. An inter-assay CV of <15% and an intra-assay CV of <10% were considered acceptable for both concentration and enrichment analysis.

#### Recovery, interferences

To evaluate analyte recovery across the sample clean-up procedure, aliquots of control pools were processed under two conditions. In the first condition, IS was added at the onset of sample preparation. In the second, IS was added only after protein precipitation and hexane extraction. Addition of IS at later stages was not feasible beyond this point because derivatization is required prior to GC–MS analysis.

Because glucose is an abundant structural isomer of fructose and its concentration varies across individuals and over time, its potential to interfere with fructose quantification was specifically examined. Control pools were fortified with high concentrations of glucose (2mM, mimicking the difference between fasted and fed plasma glucose levels). Fructose concentration and enrichment were measured in both glucose-spiked and unspiked pools to determine the extent of interference, and an observed difference of <10% (in line with intra-assay %CV) was considered acceptable.

#### Linearity

Linearity was assessed through repeated analysis of the calibration curves. The standard curve for concentration ranged from 0 nmol/ml to 1000 nmol/ml, an R^2^ of >0.998 was considered acceptable. For enrichment measurements, calibration curves were generated by dividing the integrated peak area of the tracer by that of the tracee to obtain the experimental tracer-to-tracee ratio (TTR), which was then plotted against the theoretical MPE%. Linear regression over the range from 0 to 1.5 MPE% was required to yield an R^2^ >0.998. The slope of the enrichment curve was expected to be approximately 0.01 ± 20%, confirming agreement between the experimental TTR and the theoretical MPE% values.

#### Limit of Quantification

The upper and lower limits of quantification were defined by the linear domain of the calibration curves and the linear dynamic range of the instrument. A minimum signal-to-noise ratio of >10 was used to define the lower limit of quantification (LLOQ).

#### Stability

The stability of the samples was assessed over a relevant time frame after being prepared. Multiple aliquots of the control pools were prepared, mixed and divided into vials, to ensure homogenous material for repeated injections on days 0, 2, 5, 8, 10 and 14 post-preparation. In between runs, these samples were stored at room temperature. The CV% of these results should be less than 15%, consistent with the acceptable limit for inter-assay precision. An additional experiment was performed with repeated analysis on day 33 post-preparation.

#### Carryover

Carryover was assessed using the statistical module specifically designed for carryover on the EP Evaluator software (Data Innovations LLC, Colchester, USA, Build 12.3.0.2), based on the Clinical & Laboratory Standards Institute (CLSI) EP10 guideline. For this, results of low samples measured directly after either a low or a high sample were compared. The error limit was predefined as three times the standard deviation (3×SD) of the low sample measurements.

#### Impact of anticoagulant type

The impact of anticoagulant type (EDTA versus heparine) was assessed using plasma samples collected during a clinical study (described in “clinical samples” section of the methods). At each timepoint, blood was collected in both EDTA and heparin tubes. Paired samples from both tube types were analyzed for three participants at each timepoint. To assess agreement between EDTA and heparin plasma tracer concentration, Bland-Altman analysis adapted for repeated measurements within subjects was used.

#### Effect of reduced sample volume

The effect of plasma volume taken into preparation on assay precision and accuracy was evaluated by preparing plasma aliquots of 100 µl and 50 µl, and processing them in parallel on the same day. Apart from the difference in starting material volume, the sample preparation procedure was kept identical. The experiment was repeated on a separate day to assess inter-day reproducibility.

#### Sample storage

The stability of samples under different storage conditions was evaluated by storing samples at 4 °C up to 14 days. The effect of repeated freeze–thaw cycles was also assessed for one to four cycles. Stability was calculated as the percentage compared to the freshly prepared and analyzed aliquot, and a stability of 80-120% was deemed acceptable.

#### Comparison of SIM and MRM mode

All of the above analyses were performed in MRM mode. To gain insight into the difference in performance, the experiments to assess precision and limit of quantification were performed in both SIM and MRM mode. Analyzing the same prepared samples in both acquisition modes facilitated a direct comparison, ensuring full compatibility and attributing any differences solely to the mode used.

## Results

We developed a novel GC-MS/MS method to quantify fructose concentrations and determine ^13^C_6_-fructose enrichment. Fig. 1 shows a representative chromatogram together with the mass spectra of the fructose isotopologues. We observed two chromatographic peaks (with retention times of 20.3 min and 20.8 min) corresponding to isomeric forms of derivatized fructose, both well resolved from glucose, which is also present in this plasma-derived chromatogram (Fig 1a). Despite the large physiological difference in concentration between glucose and fructose, no interference with the fructose peaks was observed. Glucose interference was further assessed in detail as part of the method validation, confirming the specificity of our method for fructose.

**Fig. 1.**
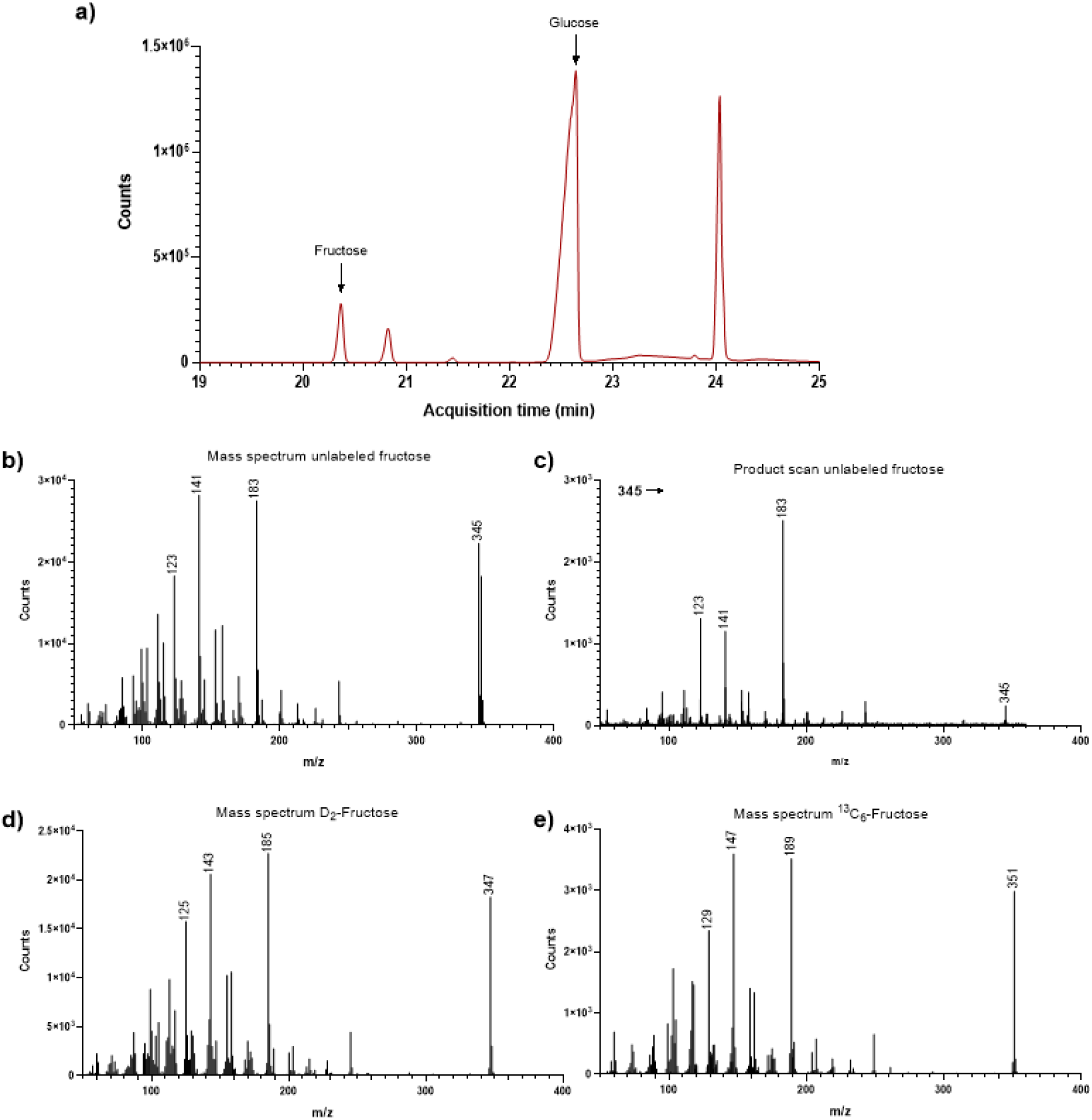
chromatogram, mass spectra and product scan of the derivatized fructose; a) Total Ion Current chromatogram recorded in MRM of a plasma sample, with transitions optimized for both fructose and glucose within a single analysis method. b, d and e) mass spectra of the isotopologues of fructose, for both D_2_-fructose and ^13^C_6_-fructose all labeled positions were preserved in the most abundant fragments; c) product scan of unlabeled fructose

The mass spectra reveal that the labeled positions are retained in the isotopologues, as reflected by the relatively abundant fragment ions at m/z 345, 347, and 351 for unlabeled fructose, D_2_-fructose, and ^13^C_6_-fructose, respectively (Fig 1b, d, e). We subsequently recorded product ion scans of these precursor ions to identify the corresponding product ions. The product ion spectrum of unlabeled fructose (Fig 1c) shows a highly abundant fragment at m/z 183, while the spectra of D_2_-fructose and ^13^C_6_-fructose show abundant fragments at m/z 185 and 189, respectively. These results confirm that the labeled positions were retained in both the precursor and product ions.

## Method validation

### Accuracy

The theoretical and observed concentrations and enrichments are presented in Table 2. The results for the blank pool were not shown, as the fructose concentration in this pool was used to calculate the theoretical concentrations of the spiked pools, and the natural background enrichment in this pool was defined as 0 MPE%.

**Table 2.** Results of the accuracy assessment. Conc: concentration; Enr: enrichment.

|  | Theoretical |  | Observed |  | Accuracy |  |
| --- | --- | --- | --- | --- | --- | --- |
|  | Conc<br>[nmol/ml] | Enr<br>[MPE%] | Conc<br>[nmol/ml] | Enr<br>[MPE%] | Conc<br>[nmol/ml] | Enr<br>[MPE%] |
| Control pool high | 598.1 | 1.44 | 554.8 | 1.30 | 93% | 90% |
| Control pool low | 133.1 | 0.43 | 128.7 | 0.40 | 97% | 94% |

The observed concentrations were obtained from duplicate analysis of each of the control pools on 5 different days. Accuracies for all parameters were within the predefined limits of 100% ± 20 %.

### Precision

Table 3 presents detailed information regarding the precision assessments. For the concentration, the intra- and inter-assay precision CVs were ≤4.6% and ≤6.5%, respectively. For the enrichment, the intra-and inter-assay precision CVs were ≤7.0% and ≤3.2%, respectively. The criteria for maximum allowable coefficients of variation as described in the materials and methods section were fulfilled.

**Table 3.** Intra- and Inter-assay precision for each of the control pools.

|  | Intra-assay precision<br>[CV%] | Inter-assay precision<br>[CV%] |
| --- | --- | --- |
| Concentration |  |  |
| - Control pool high | 3.8% | 6.1% |
| - Control pool low | 3.1% | 4.4% |
| - Control pool blank | 4.6% | 6.5% |
| Enrichment |  |  |
| - Control pool high | 1.3% | 2.7% |
| - Control pool low | 7.0% | 3.2% |

### Recovery, interferences

The analyte recovery across the sample clean-up procedure (protein precipitation and hexane extraction) was 69%. No difference in enrichment was observed during the recovery assessment.

Fortifying the plasma pools with glucose did not significantly affect the results for fructose: in control pool high, a 0.7% decrease in fructose concentration and 0.4% decrease in fructose enrichment were observed upon addition of glucose. In control pool low, an increase in fructose concentration of 3.5% and decrease in enrichment of 1.9% were observed. The concentration in the blank pool was increased by 10.6% upon addition of glucose. This was well within the predefined acceptance limit (<10%), apart from the increase in the blank pool.

### Linearity

Linearity was tested by repeated analysis of the calibration curves and complied with the validation criteria. The slope of the enrichment calibration curve was close to the theoretical value of 0.01 and within the predefined maximum allowable deviation (average slope of curves, derivatized and analyzed on five different days: 0.0108, with a precision of 3.9%.

### Limit of Quantification

The upper and lower limits of quantification were defined by the domain of linearity of the calibration curves for concentration and enrichment and the linear dynamic range of the instrument. The LLOQ was also estimated based on a signal-to-noise ratio (S/N) >10. The LLOQ for concentration was 0.1 nmol/ml. To test whether the lower limit of enrichment of the standard calibration curve (0.11 MPE%) was already approaching the true limit of linearity, we constructed a more extensive enrichment calibration curve with additional standards between the usual lowest calibration point and the unenriched standard. This confirmed that the linearity of the MRM extends further downward than the range used in the standard calibration curve (lowest enriched value: 0.11 MPE%). At a concentration of 425 nmol/ml fructose, the LLOQ for enrichment was found to be 0.02 MPE%. The resulting LLOQ values were below the lowest calibration standards used in the calibration curves.

### Stability

The stability of the prepared samples was assessed in intervals up to 14 days after preparation. The CV% of the analysis on days 0, 2, 5, 8, 10 and 14 are shown in Table 4. Additionally, a different series of samples was rerun on day 33 after preparation. The differences between the results observed on day 0 and day 33 after preparation were between 0.8% and 3.2% for concentration, and between 0.1% and 2.7% for enrichment.

**Table 4.** results of the stability test: analysis of samples on days 0 to 14 after preparation.

|  | Day 0 | Day 2 | Day 5 | Day 8 | Day 10 | Day 14 | CV% |
| --- | --- | --- | --- | --- | --- | --- | --- |
| Concentration |  |  |  |  |  |  |  |
| - Control pool high | 573.5 | 575.4 | 561.4 | 574.6 | 571.9 | 587.4 | 1.4% |
| - Control pool low | 138.1 | 134.6 | 140.4 | 138.7 | 143.1 | 143.9 | 2.4% |
| - Control pool blank | 38.3 | 38.0 | 41.3 | 40.1 | 39.8 | 41.2 | 3.5% |
| Enrichment |  |  |  |  |  |  |  |
| - Control pool high | 1.31 | 1.20 | 1.28 | 1.20 | 1.24 | 1.23 | 3.3% |
| - Control pool low | 0.46 | 0.41 | 0.41 | 0.43 | 0.42 | 0.41 | 4.1% |

### Carryover

No carryover effect was detected. The concentration of fructose observed in the high sample was 624.1 nmol/ml, and in the low sample it was 146.5 nmol. The difference between low-after-low and low-after-high was ™0.04 nmol/ml, remaining within the error limit (3xSD of low sample measurements; 15.05 nmol/ml). For the enrichment analysis, the high sample contained 1.32 MPE% ^13^C_6_-fructose, and the low sample 0.41 MPE%. The difference between low-after-low and low-after-high was ™0.02 MPE%, also within the error limit (0.11 MPE%). These results indicate the absence of carryover in both analyses.

### Impact of anticoagulant type

An example of the tracer enrichment and concentration curves observed in samples collected from peripheral plasma in either EDTA or heparin is depicted in Fig. 2a and 2b.

**Fig. 2.**
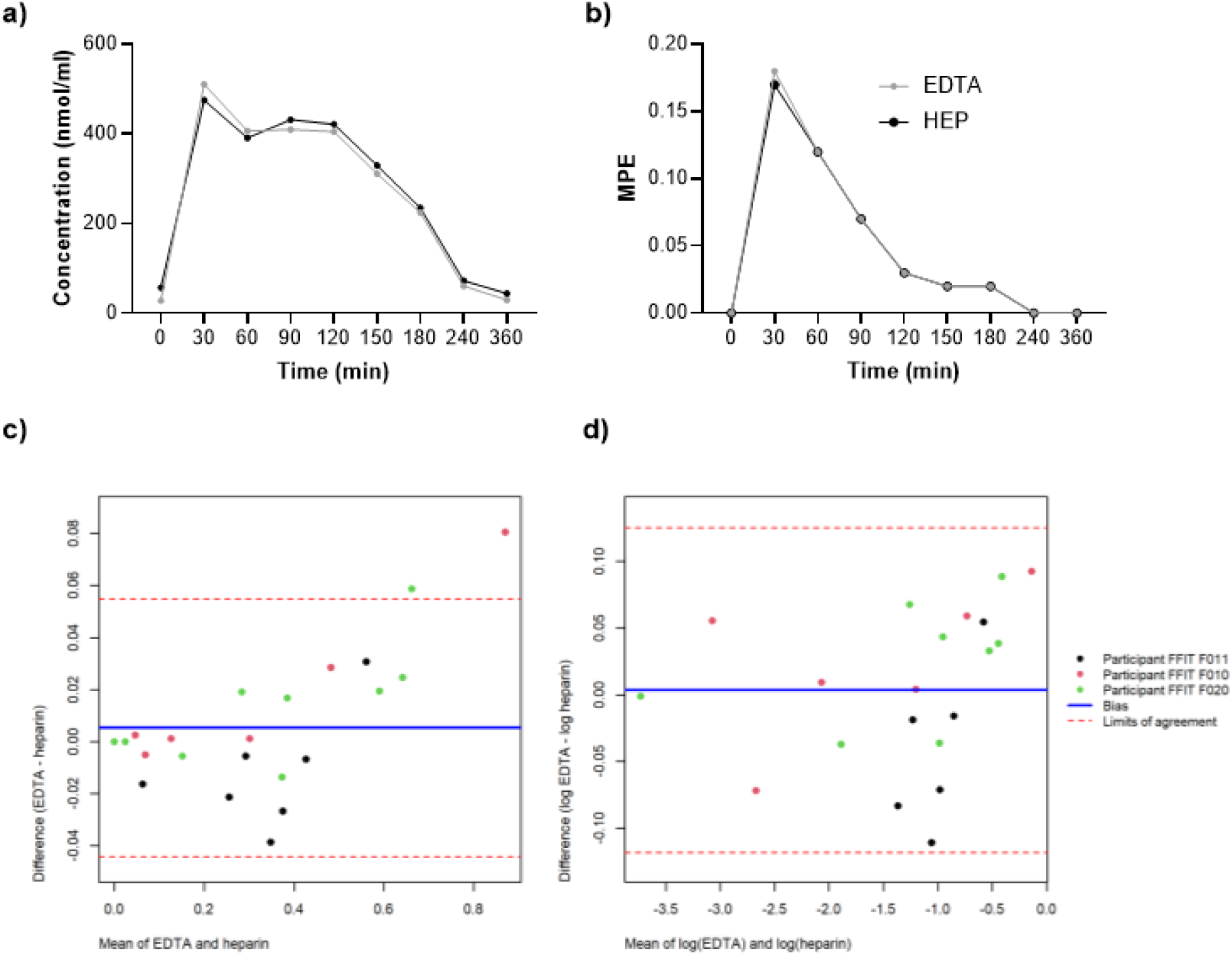
Effect of EDTA and heparin (HEP) plasma on fructose concentration and enrichment analysis. a-b) Overlay of the concentration (a) and tracer enrichment (b) curves obtained during a fructose challenge test for a single participant c-d) Bland-Altman plots comparing tracer concentration in EDTA and heparin plasma, based on 9 paired timepoints per participant. c) Raw-scale differences, showing an increase in spread with mean tracer concentration. d) Log-scale differences, expressing agreement as a relative (percentage) difference, one timepoint with a large relative difference at low absolute enrichment was excluded from the primary analysis. Solid lines represent the mean bias; dashed lines represent the 95% LoA

To assess agreement between EDTA and heparin plasma tracer concentration, Bland-Altman analysis adapted for repeated measurements within subjects was used, as each participant contributed 9 paired timepoints. On the raw scale, the spread of differences increased with the mean enrichment value (Fig. 2c), indicating that variability was proportional rather than constant across the measurement range. To test whether the differences were relative (percentage) rather than absolute, we also calculated the differences on the natural log scale. For both scales, a linear mixed model with a random intercept for participant was used to partition variance between and within subject components, from which bias and 95% limits of agreement (LoA) were derived. Timepoints where the result of either EDTA or heparin equaled zero were excluded from the log-scale analysis, as these are uninformative on a relative scale. On the log scale, one timepoint corresponding to a low absolute enrichment value in one participant showed a disproportionately large relative difference and was excluded from the primary analysis (Fig. 2d). The mean bias between EDTA and heparin was 0.003 (95% LoA: -0.118 to 0.125), corresponding to a negligible average difference and individual measurements typically differing by up to approximately 12%. Including the excluded timepoint widened the bias to -0.008 and the limits of agreement to -0.177 to 0.161 (∼15-20%), indicating that agreement was not driven by this observation. No participant or concentration dependent pattern in the differences was observed on the log scale.

### Effect of reduced sample volume

Aliquots of 100 µl and 50 µl of control samples were taken into preparation. The average CV% of duplicate analyses increased as a result of the reduced sample volume. For 100 µl aliquots the CV% was 2.1% for concentration and 1.3% for enrichment results compared to 3.5% for concentration and 1.3% for enrichment in 50 µl aliquots. Moreover, the concentration results increased with the lower sample volume: the increase was 5% for the high control, 13% for the low control, and 42% for the blank control. The observed enrichment results were less affected: a 0.2% increase for the high control, and a 2.3% decrease for the low control.

### Sample storage

The stability was calculated as percentage of the result obtained after the analysis of freshly prepared aliquot. The concentration and enrichment of the plasma samples used for this experiment were 522 nmol/ml and 1.41 MPE% for the “high” plasma sample, and 97 nmol/ml and 0.52 MPE% for the “low” plasma sample.

The stability at 4°C was acceptable for up to 14 days (average stability 101.6%, ranging from 89.4% to 116.8%), as shown in Fig. 3a. There was no significant influence of up to 4 freeze-thaw cycles (average stability 99.8%, ranging from 91.3% to 108.1%), as shown in Fig. 3b.

**Fig. 3.**
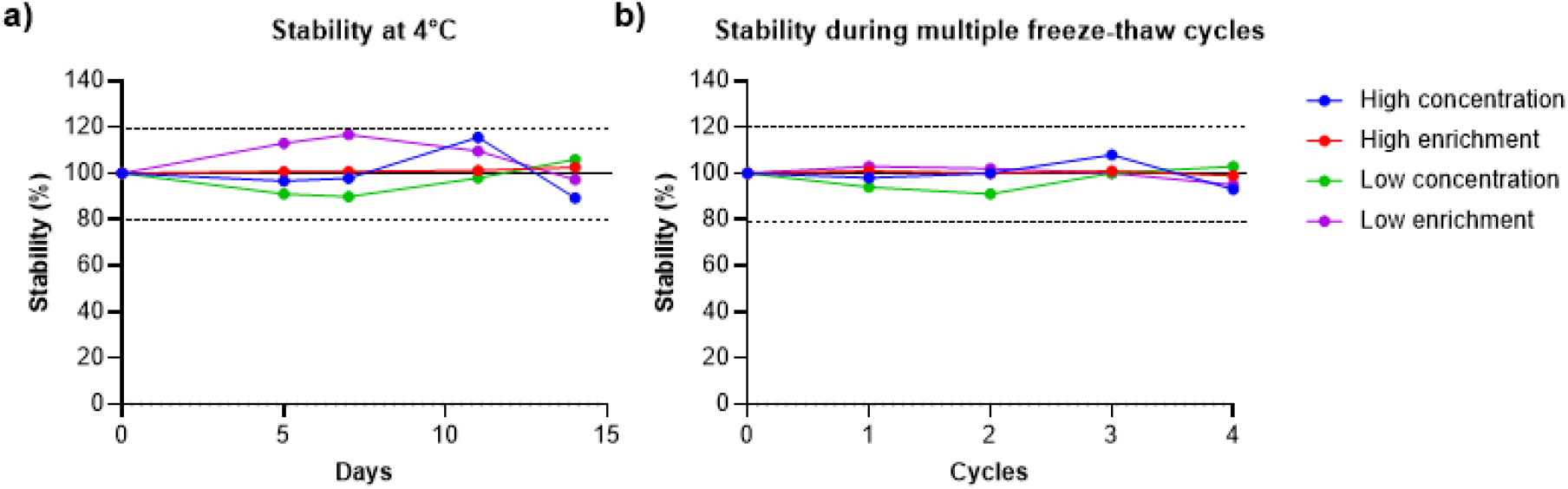
Results of the stability experiments, expressed as a percentage relative to the fresh aliquot analyzed on day 0. a) stability at 4°C up to 14 days and b) during multiple freeze-thaw cycles. The solid line at 100% represents the nominal value (no change relative to day 0), and the dashed lines at 80% and 120% indicate the predefined acceptance limits

### Comparison of SIM and MRM mode

The same prepared samples used to determine precision and the limit of quantification were analyzed in both SIM and MRM mode, allowing direct comparison of the two acquisition modes. The precision results are presented in Table 5. The relative difference between duplicate injections was found to be 1.3% for concentration and 5.3% for enrichment in MRM mode, and 1.5% for concentration and 3.6% for enrichment in SIM mode. The extensive enrichment calibration curve analyzed in both acquisition modes was depicted in Fig. 4. The difference in offset between the calibration curves obtained in the two different modes is attributable to the larger fragment monitored in SIM, which contains more atoms and therefore has a higher probability of containing a naturally occurring stable isotope; MRM monitors a smaller fragment, resulting in lower natural enrichment and, consequently, a smaller offset.

**Table 5.** Intra- and Inter-assay precision for each of the control pools, comparison of performance of MRM and SIM.

|  | Intra-assay precision [CV%]<br>MRM vs SIM | Inter-assay precision [CV%]<br>MRM vs SIM |
| --- | --- | --- |
| Concentration |  |  |
| - Control pool high | 3.8% vs 3.6% | 6.1% vs 4.7% |
| - Control pool low | 3.1% vs 2.8% | 4.4% vs 4.3% |
| - Control pool blank | 4.6% vs 5.1% | 6.5% vs 9.6% |
| Enrichment |  |  |
| - Control pool high | 1.3% vs 0.9% | 2.7% vs 7.3% |
| - Control pool low | 7.0% vs 6.2% | 3.2% vs 17.5% |

**Fig. 4.**
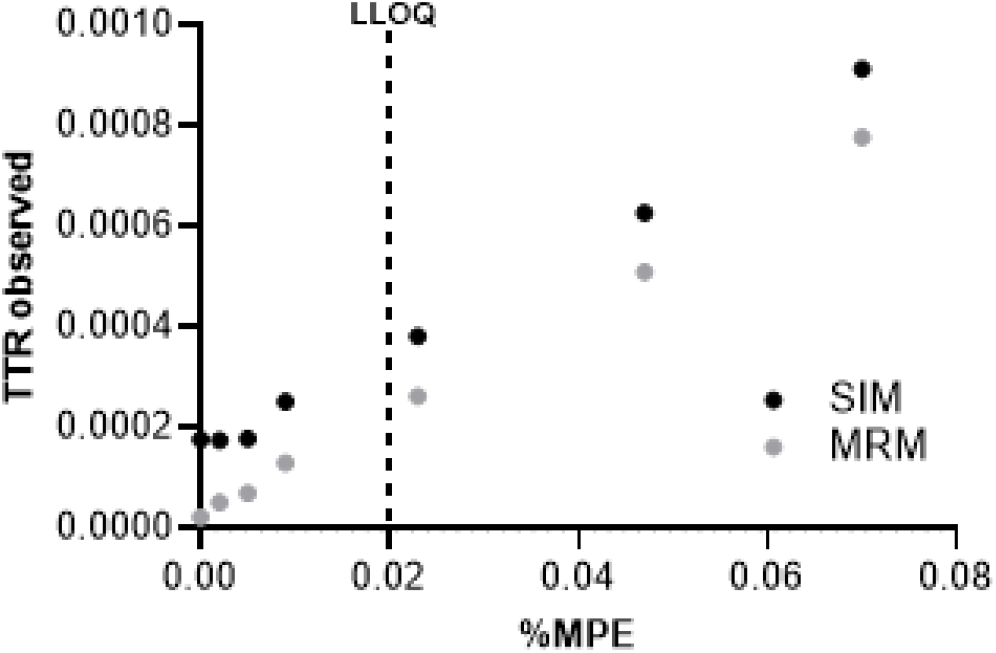
Extended enrichment calibration curve, showing results for both SIM and MRM. The dashed line indicates the LLOQ determined for MRM. The TTR observed was calculated from the area of the ^13^C_6_-fructose peak divided by the area of the unlabeled fructose peak observed in the standards of the calibration curve.

## Discussion

High dietary fructose consumption is a key driver of obesity and related cardiometabolic diseases, underscoring the importance of studying fructose metabolism in humans. Stable isotope tracers, such as ^13^C_6_-fructose, are essential for this purpose, but require a highly sensitive and validated analytical method capable of accurately quantifying fructose concentrations and isotopic enrichment. Here we present a validated method that specifically and accurately quantifies both unlabeled and ^13^C_6_-fructose. Moreover, the measurements have an inter and intra assay precision below 10% with no detectable carryover, and a LLOQ of 0.1 nmol/ml for concentration and 0.02 MPE% for enrichment.

### Method rationale and specificity

A key objective was to determine whether the derivatization procedure routinely applied to glucose could also be used for fructose, allowing both analytes to be measured independently in the same prepared sample after appropriate dilution. This raised the possibility that glucose, present at much higher physiological concentrations (millimolar range), could interfere with fructose quantification (micromolar range). This was addressed during validation by achieving chromatographic separation of fructose and glucose (Fig. 1a), and by confirming no detectable effect on fructose quantification in clinical samples containing physiological glucose concentrations (5–10 mM), with accuracy remaining high under these conditions. These findings indicate that co-elution or ion suppression from glucose does not compromise fructose analysis, supporting the use of a single sample preparation for both analytes. This is in line with a previously reported GC-MS (SIM) method using a similar (though not identical) derivatization strategy, which likewise allowed separate quantification of fructose and glucose (36).

### Rationale for transition selection

Using a tracer in which all labeled positions are retained in both the precursor and product ions of the MRM transition minimizes the risk that unlabeled isotopologues or in-source artifacts contribute to the measured signal. We have previously reported a method in which the product ion instead corresponded to a fragment of the derivatizing group, which also performed well in increasing the sensitivity of the method (35); the present approach adds a further layer of specificity by maximizing retention of labeled carbons in the monitored transition.

### Is MRM strictly necessary?

MRM provides increased selectivity relative to SIM, because the probability that a co-eluting compound shares both the Q1 precursor ion mass and the product ion mass is low. This selectivity translates into higher sensitivity, consistent with our own observations and prior reports, e.g. (35, 37), and is particularly advantageous at low isotopic enrichment or low analyte concentration. Accordingly, the increased sensitivity of the present MRM method for concentration analysis is reflected in a lower LLOQ compared to a previously reported method employing a similar derivatization strategy, but analyzed in SIM mode (36).

The increased sensitivity of enrichment analysis in MRM mode relative to SIM mode is illustrated in Fig. 4, which compares SIM and MRM calibration curves for low enrichments: the MRM curve shows improved linearity, with a lower intercept reflecting a reduced contribution from naturally occurring isotopes in the smaller product ion. More generally, when low enrichments are expected, a more heavily labeled tracer should be chosen as the natural isotopic abundance produces a lower background enrichment, that is substantial relative to the true tracer signal for e.g. singly labeled metabolites. As the number of labeled positions increases, this background contribution decreases, improving precision at low enrichment. However, MRM is not universally advantageous: for example for analytes whose EI fragments are already small, further fragmentation can yield product ions that are non-specific or of low abundance (38). Is MRM therefore strictly necessary? In the present study, MRM performed better than SIM at the low concentrations and enrichments measured in clinical plasma samples, and is therefore preferred where GC-QQQ instrumentation is available. Acceptable results can nonetheless be obtained using SIM, which is relevant for laboratories without access to triple-quadrupole instrumentation. For such laboratories, an alternative approach for improving precision is to increase the tracer dose, at the cost of higher trial expense.

### Recovery, accuracy, and precision

Method validation confirmed acceptable recovery. Although a fraction of fructose was lost during sample clean-up, this loss did not affect the reported isotopologue ratios, since the labeled internal standard was added prior to sample preparation and isotopologues ratios are preserved despite incomplete recovery. Accuracy was consequently high, and both intra- and inter-assay precision were excellent.

### Linearity and dynamic range

Linear calibration curves were obtained for both concentration and enrichment. At high concentrations, curves occasionally showed downward curvature attributable to detector saturation rather than peak overloading (peak shape remained Gaussian). Saturation, occurring predominantly at the signal of unlabeled fructose, results in underestimated concentration or overestimated enrichment, this is corrected by further diluting the final extract in ethyl acetate. Because the linear dynamic range is instrument-specific, it should be established for each system by preparing a concentrated sample or standard together with a corresponding dilution series in ethyl acetate. The range where isotope ratios remain constant defines the working dynamic range for analysis to avoid biased results. Given the wide concentration range expected among clinical study samples, any standards or samples exceeding the linear range should be re-analyzed at reduced injection volume or after dilution.

### Method performance

The LLOQ was well below the lowest calibration standard for both concentration and enrichment, and well below the lowest fructose concentration observed in plasma samples to date. Reduced sample volume did not compromise overall precision, although a concentration dependent deviation was observed. To maximize comparability and reliability across samples, especially low-concentration samples such as those obtained after fasting, we recommend taking a standardized 100 µL aliquot into preparation.

### Stability and matrix flexibility

Prepared samples were highly stable, permitting repeated analysis when re-dilution is required due to exceedance of the linear range or to instrument downtime. Results were unaffected by extended sample storage and freeze-thaw cycles, tested under conditions realistic for typical clinical study logistics. Because clinical samples are often irreplaceable, the ability to measure fructose reliably after these conditions makes this protocol particularly well suited for patient sample analysis.

### Consideration for plasma collection

EDTA and heparin plasma produced comparable results and can likely be used interchangeably for fructose tracer analysis. Bland-Altman analysis showed negligible systematic bias between EDTA and heparin plasma for tracer enrichment measurements, with individual measurements agreeing within approximately 12–20%, depending on whether one low-concentration timepoint was included. This imprecision could be due to the amplification of the assay variation at low concentrations rather than a true anticoagulant-dependent effect. Confirmation in a larger sample would strengthen confidence in the observed agreement. As best practice, we nonetheless recommend using a single anticoagulant consistently within a given clinical study.

Here we describe a validated method to measure the concentration and enrichment of fructose, further tested in a clinical setting using peripheral plasma samples after an FCT. In this FCT, subjects orally ingest an unlabeled fructose load together with a ^13^C_6_-labeled tracer to investigate whole-body fructose catabolism under various (disease) conditions. The samples used for this validation served as a physiological example to test the protocol, as it is intended for further application in clinical trials investigating altered fructose handling following different treatments. The FFIT study (Dutch Clinical Trial Register, registration ID: NL-OMON24239) aims to elucidate the effects of fecal microbiota transplantation (FMT) on microbial fructose fermentation in the intestine, by comparing healthy donor FMT with autologous FMT in patients with metabolic syndrome. Furthermore, our group has several ongoing randomized controlled trials examining the effects of high-versus isocaloric low-fructose diets on fructose whole-body catabolism and its clinical effects in individuals with morbid obesity (Dutch Clinical Trial Register, NL-OMON51827) and type 2 diabetes (NL-OMON51729). This highlights the relevance of establishing an accurate fructose tracing method for advancing the understanding of fructose catabolism and its role in obesity and related metabolic disease.

## Conclusions

We optimized and validated a method for the analysis of fructose enrichments and concentrations at low levels. Oximation and per-acetate derivatives were suitable for this analysis and stable over time. We demonstrate that MRM improves method performance only at low concentrations and enrichments relative to SIM. This method can be applied in stable-isotope tracing studies to investigate fructose metabolism in depth and to further understanding of the role of fructose in the development of obesity and related cardiometabolic diseases.

## Statements and declarations

Plasma samples were acquired, following informed consent, from adult participants of a clinical study (Dutch Clinical Trial Register, registration ID: NL-OMON24239). The study was conducted in accordance with the ethical standards and, where required, approved by the ethics committee of the Amsterdam UMC.

Partial financial support was received from the Academic Education and Research sector plans of the Dutch Ministry of Education, Culture and Science for the appointment of D.v H.. M.N. is supported by a personal NWO VICI grant 2020 (09150182010020) and an ERC Advanced grant (101141346) on which M.R.M. and F.H.M.W. are appointed. M.N. is co-founder and member of the Scientific Advisory Board of Caelus Pharmaceuticals and Advanced Microbiota Therapeutics (Both Spinouts of AmsterdamUMC), Amsterdam the Netherlands. However, none of these are directly relevant to the current paper. There are no patents, products in development or marketed products to declare. F.M.V. has acted as a consultant to Scenic Biotech and Spolia Therapeutics; is a member of the Danone-funded scientific advisory board of the European Metabolic Group; is an inventor on patent US11650212B2 (a newborn screening method for cerebrotendinous xanthomatosis), assigned to Amsterdam UMC Foundation (formerly Academic Medical Center); and heads a fee-for-service core facility that performs contract research for commercial parties and collaborates with mass spectrometry manufacturers including Bruker. No other financial or non-financial interests that could be perceived to influence the work reported here are declared. The other authors declare no conflict of interest.

## Data availability statement

Data are available from the authors by request.

## Abbreviations

T2D: type 2 diabetes
MASLD: metabolic dysfunction-associated steatotic liver disease
DNL: de novo lipogenesis
GC-(MS/)MS: gas chromatography-(tandem) mass spectrometry
GC-QQQ: gas chromatography–triple quadrupole mass spectrometry
MRM: multiple reaction monitoring
SIM: selected ion monitoring
IS: internal standard
MW: molecular weight
m/z: mass to charge ratio
CV: Coefficient of variation
CLSI: Clinical & Laboratory Standards Institute
LLOQ: lower limit of quantification
S/N: signal-to-noise ratio
MPE%: molar percent excess
LoA: limits of agreement
RCTs: randomized controlled trials
FMT: fecal microbiota transplantation

## Acknowledgements

We would like to thank the participants for their contribution to the clinical study and Amrita Tjaris – Eeftinck Schattenkerk for technical assistance.

## Author contributions

All authors contributed to the study conception and design. Material preparation, data collection and analysis were performed by Melany Rios Morales, Florine Westerbeke and Dewi van Harskamp. The first draft of the manuscript was written by Melany Rios Morales, Florine Westerbeke and Dewi van Harskamp and all authors commented on previous versions of the manuscript. All authors read and approved the final manuscript.

